# Epigenetic editing of Cartpt promotes acquisition and extinction of cocaine memory

**DOI:** 10.1101/2025.11.20.689329

**Authors:** Julia J. Winter, Marisol Hooks, Keegan S. Krick, Alexandra Goldhamer, Ronald W. DiTullio, Kyle S. Czarnecki, Claude-Ericka Ekobeni, Chloe Han, Kiara L. Rodríguez-Acevedo, Brandon W. Hughes, Molly Estill, Collin D. Teague, Aarthi W. Ramakrishnan, Li Shen, Eric J. Nestler, Elizabeth A. Heller

## Abstract

In classic disease models, removing a pathological insult restores homeostasis. Yet, addiction persists far beyond the period of active drug use. Cocaine abstinence induces changes in gene expression and neuronal signaling in reward-related brain regions that limit recovery during abstinence. We found that 2 weeks of abstinence increased *Cartpt* (cocaine- and amphetamine-regulated transcript) in the mouse nucleus accumbens and decreased repressive H3K27me3 at the *Cartpt* locus. While endogenous CART peptide is best described for its anorexigenic function, it is also implicated in human addiction and dopamine homeostasis. To test the causal relevance of *Cartpt* chromatin remodeling, we used CRISPR-based epigenetic editing tools, dCas9-FOG1 and dCas9-JMJC-ZF, to manipulate H3K27me3 at *Cartpt in vivo*. Enriching H3K27me3 in D1 neurons repressed *Cartpt* expression and augmented acquisition and extinction of cocaine preference. These results show that CRISPR epigenetic editing can recapitulate endogenous chromatin states to modulate addiction-related behavior, highlighting broad therapeutic potential of both *Cartpt* and epigenetic editing.

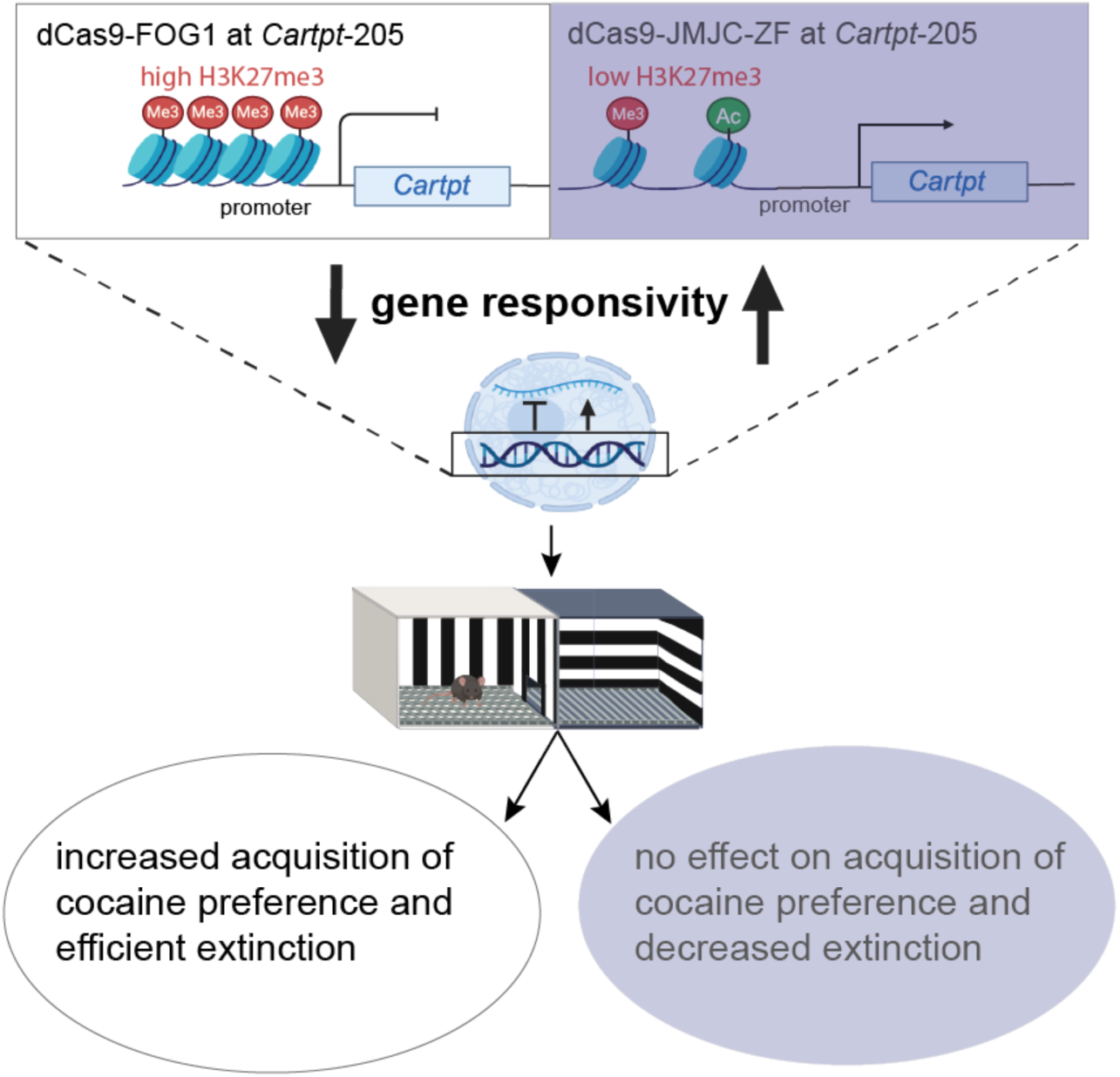

## INTRODUCTION

Cocaine addiction, a prevalent and treatment-resistant neuropsychiatric condition, remains a global public health crisis, affecting millions of individuals and placing a substantial burden on healthcare systems^1^. The persistence of addiction extends far beyond the period of active drug use, as it triggers long-lasting neurobiological changes that limit recovery during abstinence^2,3^. Understanding the mechanisms behind these enduring adaptations is critical for developing effective treatments aimed at facilitating recovery and preventing relapse. The persistence of cocaine-induced neuroadaptations involves lasting changes to gene expression in brain regions within the mesolimbic dopamine circuit^4,5^. Many studies focus primarily on the nucleus accumbens (NAc)^6–8^, which plays a key role in regulating cocaine-reward behavior by responding directly to dopamine signaling. Cocaine increases dopamine in the NAc, triggering downstream epigenetic regulation of gene expression^9,10^. Unbiased transcriptomic analyses in the NAc following cocaine exposure reveal that the number of differentially expressed genes that increase over time, with 2-7x more genes uniquely regulated at 1 month of abstinence compared to 1 day^5,11^. We hypothesize that the interaction between cocaine and the epigenetic state of a gene plays a critical role in modulating the gene’s responsivity to cocaine-mediated signaling^9,12^. Mechanistically, epigenetic editing in the brain has gained considerable attention for its promise in rare neurological disorders. Recent studies have demonstrated that targeted epigenetic modulation can restore gene expression and improve cellular and behavioral outcomes in models of neurodevelopmental disorders including Fragile X syndrome^13^, prion disease^14^, and in highly prevalent psychiatric disorders^15–17^. These advances underscore the potential of epigenetic editing to reprogram disease-relevant gene regulation. Understanding these mechanisms can help restore neuronal function after cocaine-induced changes and may offer new therapeutic avenues.

Here, we present new insights into the specific epigenetic mechanism of cocaine- and amphetamine-regulated transcript peptide (*Cartpt*) in cocaine reward. *Cartpt* is an important molecule in cocaine reward behavior that is persistently activated during cocaine abstinence^11^. In addition to our published findings, the literature supports the relevance of *Cartpt* to cocaine addiction^18^. Analysis of human post-mortem brains finds an upregulation of CARTPT mRNA in the NAc, a finding replicated in rodent models following acute cocaine administration^6,19,20^. Injecting CART peptides into the NAc reduces cocaine-induced locomotion and inhibits the rewarding effects of cocaine self-administration^21–25^. While the precise mechanism is unclear, studies indicate that CART in the NAc responds to increased extracellular dopamine levels and inhibits dopaminergic transmission^26–28^. Outside of cocaine reward, CART peptide also plays an important role in alcohol use disorder and other psychiatric disorders including anxiety^29^. Furthermore, in the context of alcohol use disorder CART peptide exhibits a sexually dimorphic effect with female CART knockout mice showing decreased alcohol intake compared to control whereas male CART knockout mice increased alcohol intake^30^.

Prior studies find that *Cartpt* expression after cocaine exposure is dynamic. Specifically, *Cartpt* is activated immediately after cocaine exposure in a cell type-specific manner^31^, returns to baseline after 1 day of abstinence and is then re-activated at 1 month of abstinence in the NAc^11^. The current study further refines these data, finding that *Cartpt* is activated at 2 weeks, but not at 4 or 7 days of cocaine abstinence. To interpret these patterns of gene activation, we referred to our prior study that found delayed *Cartpt* activation is associated with the depletion of the repressive histone posttranslational modification histone 3 lysine 27 trimethylation (H3K27me3) and the enrichment of the activating mark histone 3 lysine 27 acetylation (H3K27ac) at the *Cartpt* promoter^11^. This aligns with various studies showing that cocaine persistently remodels H3K27me3 landscape in brain reward regions, which facilitates aberrant gene activation linked to drug sensitization, synaptic plasticity, and relapse vulnerability^10,32–34^.

To characterize the direct causal relevance of H3K27me3 to delayed, abstinence-driven *Cartpt* expression and its behavioral role, we applied novel CRISPR-based epigenetic editing in the mouse NAc using dCas9-FOG1^35^ and dCas9-JMJC-ZF. We found that H3K27me3 enrichment at *Cartpt* is sufficient to alter acquisition and extinction of cocaine-context associations. CRISPR-based epigenetic editing is the only approach to determine the causal relevance of cocaine-associated chromatin changes and harbors the potential to bring targeted epigenetic therapies in humans closer to clinical reality.

## RESULTS

### Prolonged cocaine abstinence regulates expression of behaviorally relevant genes, including *Cartpt*

To investigate transcriptomic and chromatin changes across abstinence, we treated mice with chronic cocaine for 10 days and assessed gene expression in the NAc at 4, 7, and 14 days of abstinence (Fig. 1a). We observed a progressive increase in the number of differentially regulated genes (DEGs) across cocaine abstinence (Fig. 1a). *Cartpt* mRNA was increased only after 14 days of abstinence (Fig. 1a), which we confirmed by qPCR (Fig. 1b). Gene ontology (GO) analysis showed predominantly regulation of ion channel activity and transmembrane transport (Fig. 1c). We observed enrichment for behaviorally relevant terms only after 14 days of abstinence (Fig. 1d). By 14 days of abstinence, cognition emerged as the top functional category of all DEGs combined (Fig. 1d, left panel, purple). GO enrichment analysis of upregulated genes revealed significant overrepresentation of processes related to axonogenesis, cognition, and development (Fig. 1d, middle panel, red). In contrast, downregulated genes were enriched for behaviorally relevant terms, including locomotion (Fig. 1d, right panel, blue). Additionally, we found that *Cartpt* contributed to, among others, feeding behavior, positive regulation of MAPK pathway, and neuropeptide signaling pathway at 14 days of abstinence.

**Figure 1.**
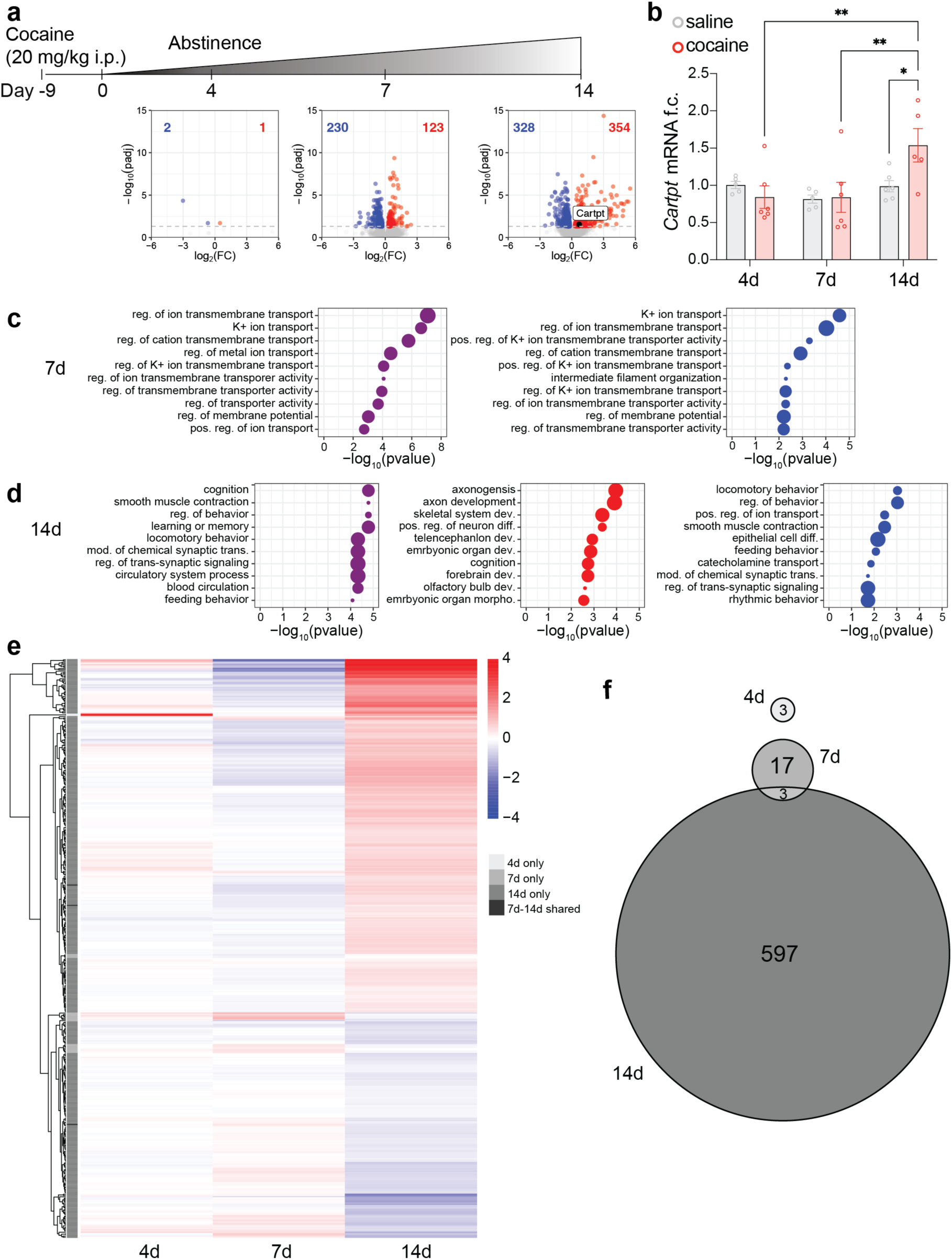
Transcriptional profiling across cocaine abstinence finds delayed enrichment of behavioral functions, including expression of *Cartpt*. **a** Schematic of i.p. cocaine injections (20 mg/kg) and abstinence time course. Volcano plots of differentially expressed genes (DEGs), individual DESeq2 objects were analyzed for each timepoint using ‘treatment’ as a variable, left to right: 4 days of abstinence, 7 days of abstinence, 14 days of abstinence. Blue indicates significantly downregulated genes, red indicates significantly upregulated genes, p<0.05 n=4. **b** qRT-PCR analysis confirmed significant increase of *Cartpt* mRNA expression compared to saline-treated mice as well as all other cocaine abstinence timepoints. Mixed-effects analysis, significant effect of timepoint, p=0.0165 and trending effect of timepoint x treatment, p=0.0600, followed by Tukey’s multiple comparisons test, 14d abstinence saline vs cocaine *p=0.0139, cocaine 4d abstinence vs 14d abstinence **p=0.0069, cocaine 7d abstinence vs 14d abstinence **p=0.0067, n-number for all treatments and groups n=5-6. **c** Gene ontology analysis of DEGs at 7 days of abstinence, p<0.05, term size=1000, purple dots represent all changes regardless of directionality, blue dots represent GO analysis on downregulated genes only. **d** Gene ontology analysis of DEGs at 14 days of abstinence, p<0.05, term size=1000, purple dots represent all changes regardless of directionality, red and blue dots represent GO analysis on upregulated and downregulated genes separately. **e** For comparisons over time, data was combined in one DESeq2 object and analyzed using ‘treatment’, ‘timepoint’ and ‘treatment x timepoint’ as a variable. Heatmap of DEG expression levels at 14 days, with expression patterns shown for each timepoint in the first and second column. Genes are hierarchically clustered by expression profile at 14 days. Comparisons are shown at 4, 7, and 14 days cocaine vs saline control, padj < 0.05, no Log2FC cutoff. On the left, the grey sidebar indicates gene overlap between timepoints, light to dark grey shows 4-, 7-, and 14-day-DEGs, black indicates overlapping genes between 7 and 14 days. **f** Venn diagram showing the overlap of DEGs across all abstinence time points. Each circle represents the set of DEGs (padj < 0.05) identified at a given time point. Overlapping regions indicate genes shared between two or more time points, while non-overlapping regions represent time point–specific DEGs. The largest number of DEGs was observed at 14 days, with minimal overlap between time points, indicating largely distinct transcriptional responses over time.

To directly compare the identity, number, and magnitude of differential gene expression over time, we expanded the DESeq2 model to include timepoint as a variable (Fig. 1e). A clear increase in the number of DEGs was observed only at 14 days of abstinence. This increase included a greater number of both upregulated and downregulated genes, as well as higher absolute log2 fold changes compared to 4 and 7 days. The identity of the DEGs also differed, with limited overlap between time points (Fig. 1e), indicating a distinct transcriptional profile at 14 days. Notably, the changes observed at 4 days were absent at 7 and 14 days of abstinence (Fig. 1f).

### CRISPR epigenetic editing of *Cartpt* H3K27me3 via dCas9-FOG1 and dCas9-JMJC-ZF bidirectionally regulates *Cartpt* expression in neurons *in vivo*

We next defined the direct causal relevance of H3K27me3 to *Cartpt* expression and cocaine reward using *in vivo* CRISPR epigenetic editing. First, we analyzed H3K27me3 landscape of *Cartpt* in the NAc using CUT&RUN. We found broad H3K27me3 enrichment at *Cartpt*, confirming our prior cell-type-specific analysis in D1 and D2 medium spiny neurons (MSNs) in the NAc^36^ (Fig. 2a). We designed a *Cartpt* single-guide RNA (sgRNA) that binds 205 base pairs upstream of the *Cartpt* transcription start site (TSS) at a locus of endogenous H3K27me3 enrichment (Fig. 2a and b). We confirmed the efficacy of *Cartpt*-sgRNA for CRISPR-based manipulation *in vitro* by co-expression with dCas9-VP64 activator in N2a cells (Fig. 2c). Next, we applied CRISPR-dCas9-FOG1 (Friend of GATA1) (Fig. 2d), which drives H3K27me3 enrichment in HCT116 cells^35,37^. We co-transfected N2a cells with dCas9-FOG1 and *Cartpt*-sgRNA and found decreased *Cartpt* mRNA expression (Fig. 2e). For intra-NAc CRISPR, we cloned dCas9-FOG1 into a neuron-specific expression vector under the control of the hSynapsin1 promoter and expressed this in NAc with either the *Cartpt*-sgRNA or control-sgRNA, ipsilaterally (Fig. 2d and f, Fig. S2a). We confirmed NAc-specific plasmid expression in fresh coronal sections using GFP visualization (Fig. 2f) and transfection efficiency via qPCR analysis 4 days post-surgery (Fig. S2b-d). CRISPR-dCas9-FOG1 and *Cartpt*-sgRNA enriched *Cartpt* H3K27me3 and reduced *Cartpt* in NAc compared to control-sgRNA on experimental days 4 and 7 in male and female mice (Fig. 2g and j, Fig. S2e and f). This effect was transient as *Cartpt* levels returned to baseline 28 days after injections (Fig. 2h). Targeting CRISPR to a promoter region can sterically interfere with transcription initiation^38^. To confirm that *Cartpt* repression was produced by the FOG1 domain rather than non-catalytic mechanisms we compared dCas9-FOG1 to dCas9 fused only to a synthetic affinity tag, AMtag^11,39^. Co-transfection of dCas9-FOG1 and *Cartpt*-sgRNA repressed *Cartpt* mRNA levels, relative to co-transfection of dCas9-AMtag and *Cartpt*-sgRNA (Fig. 2i). We next measured H3K27me3 using semiquantitative chromatin immunoprecipitation (ChIP-qPCR) of H3K27me3 on day 4 post-injection of dCas9-FOG1 and *Cartpt*-sgRNA using qPCR primers (G1-G3) spanning 512 bp centered on the *Cartpt*-sgRNA (*Cartpt* TSS -205) (Fig. 2l). We found that *Cartpt* H3K27me3 was enriched at primer G2 when dCas9-FOG1 was transfected with *Cartpt*-sgRNA, relative to control-sgRNA (*trending*, p<0.1, Fig. 2j, Fig. S2g and h) and confirmed that H3K27me3 enrichment correlated to dCas9-FOG1 expression level when co-injected with *Cartpt*-sgRNA but not control-sgRNA (Fig. 2k).

**Figure 2.**
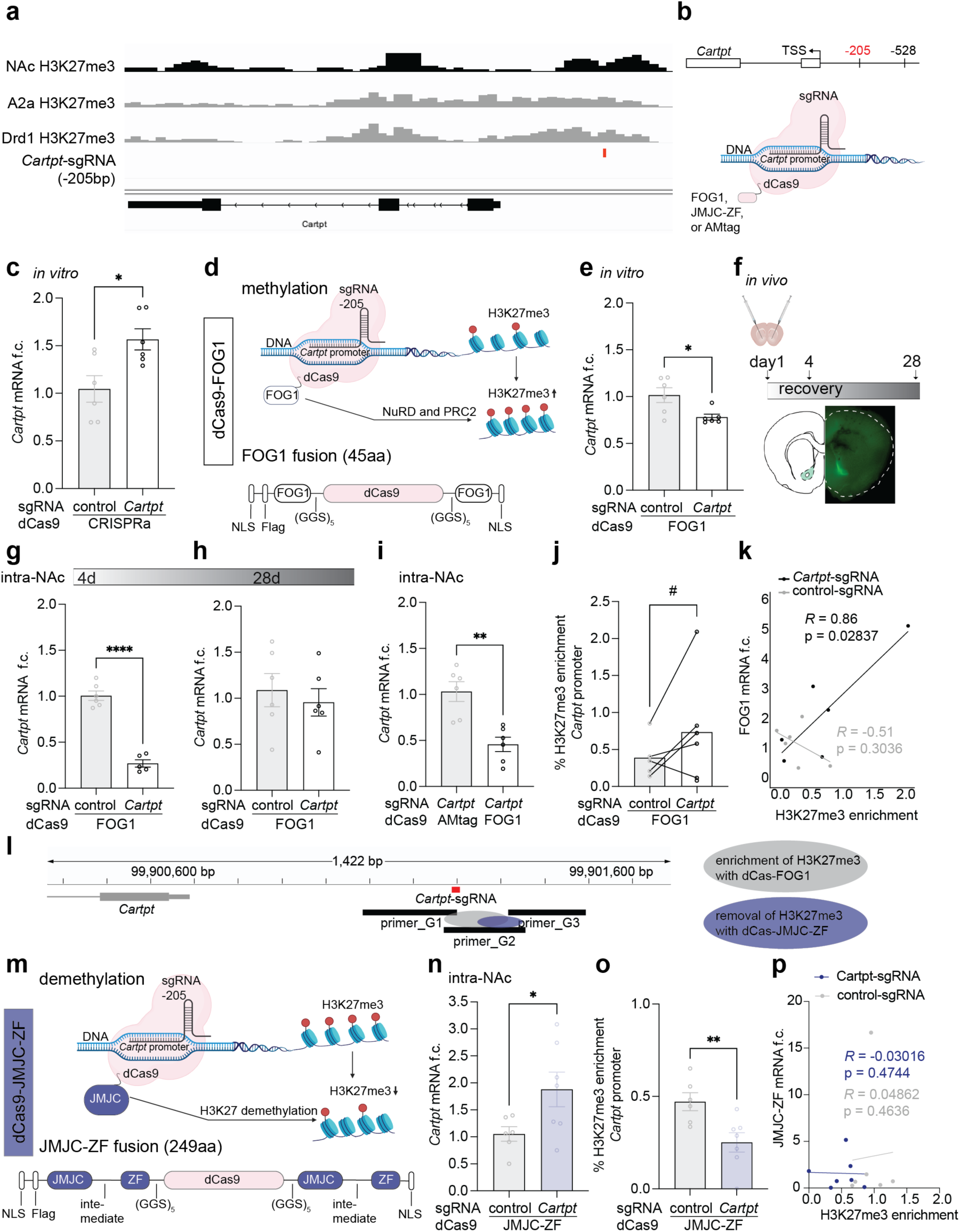
Bidirectional epigenetic editing of H3K27me3 at the *Cartpt* promoter. **a** IGV genome browser view of the *Cartpt* locus for different H3K27me3 chromatin profiling methods and cell types. From top to bottom: NAc bulk CUT&RUN data from animals treated with cocaine for 10 days, tissue was taken after 4 days of abstinence, A2a MSN and Drd1 MSN H3K27me3 ICuRuS from Carpenter et al., 2022^36^, location of the *Cartpt*-205 sgRNA is indicated in red. **b** Schematic representation of *Cartpt*-sgRNA location (-205 bp from TSS). **c** dCas9-VP64 and *Cartpt*-sgRNA increased *Cartpt* mRNA in N2a cells (unpaired two-tailed t test with Welch’s correction, *p=0.0160, t=2.924, df=9.517, sgRNA-205, n=6), compared to control-sgRNA. **d** Schematic of dCas9-FOG1 methylation mechanism at the *Cartpt* promoter. **e** dCas9-FOG1 and *Cartpt*-sgRNA decreased *Cartpt* mRNA in N2a cells (Mann Whitney test, two-tailed, *p=0.0260, Mann-Whitney U=4, sgRNA-528, n=6), compared to dCas9-FOG1 and control-sgRNA. **f** Experimental time course, including intracranial injection of CRISPR tools and brain tissue collection (top). Fluorescent microscopy confirms localization of CRISPR plasmids to NAc at day 4 (bottom). **g** *Cartpt* mRNA is reduced in NAc at day 4 after injection of dCas9-FOG1 and *Cartpt*-sgRNA, compared to control-sgRNA (unpaired two-tailed t test with Welch’s correction, ****p<0.0001, t=11.25, df=8.847, n=5-6 mice/group). **h** *Cartpt* mRNA f.c. was unchanged 28 days after injection of dCas9-FOG1 with either control-sgRNA or *Cartpt*-sgRNA. Unpaired two-tailed t test with Welch’s correction, p=0.5845, t=0.5657, df=9.670, n=6. **i** dCas9-FOG1 and *Cartpt*-sgRNA reduced *Cartpt* mRNA *in vivo* relative to dCas9-AMtag/*Cartpt*-sgRNA (unpaired two-tailed t test with Welch’s correction, **p=0.0019, t=4.319, df=9.119, n=6 mice/group). **j** dCas9-FOG1/*Cartpt*-sgRNA enriched H3K27me3 at the *Cartpt* promoter with a trending effect when assessed by primer_G2 (paired t test, one-tailed, #p=0.0618, t=1.945, df=4, n=5-6 mice/group), relative to dCas9-FOG1/control-sgRNA. **k** Correlation between H3K27me3 enrichment and dCas9-FOG1 mRNA, Pearson correlation, each point represents an individual sample, linear regression lines are shown for each condition, n=5-6. **l** Schematic representation of qChIP-qPCR primer pairs (G1-G3) relative to *Cartpt* gene locus, grey (FOG1) and purple (JMJC-ZF) circles indicate regions of epigenetic editing. **m** Schematic of dCas9-JMJC-ZF demethylation mechanism at the *Cartpt* promoter. **n** dCas9-JMJC-ZF and *Cartpt*-sgRNA increased *Cartpt* mRNA *in vivo* (unpaired two-tailed t test witch Welch’s correction, *p=0.0440, t=2.387, df=8.006, n=6-7 mice/group) compared to control-sgRNA. **o** dCas9-JMJC-ZF/*Cartpt*-sgRNA de-enriched H3K27me3 at the *Cartpt* promoter when assessed by primer_G2 (unpaired t test with Welch’s correction, **p=0.0052, t=3.085, df=11.00, n=6 mice/group), relative to dCas9-JMJC-ZF/control-sgRNA. **p** Correlation between H3K27me3 enrichment and *Cartpt* mRNA, Pearson correlation, each point represents an individual sample, linear regression lines are shown for each condition, n=6.

To achieve bidirectional regulation of *Cartpt* we developed a new epigenetic editing tool, dCas9-JMJC-ZF, for the targeted demethylation of H3K27me3 (Fig. 2m, Fig. S2a). We identified a 249 amino acid fragment of mouse JMJC as the effector domain to demethylate H3K27me3 and a zinc finger (ZF) motif downstream of JMJC to specify K27 targeting^40,41^. We tethered the JMJC and ZFs to the N- and C-termini of dCas9 to generate a specific epigenetic editing tool for the removal of H3K27me3. *Cartpt* mRNA was increased in mice 4 days after intra-NAc injections of dCas9-JMJC-ZF and *Cartpt*-sgRNA relative to dCas9-JMJC-ZF and control-sgRNA (Fig. 2n). We confirmed that dCas9-JMJC-ZF reduced H3K27me3 at the *Cartpt* locus using ChIP-qPCR only at the area detectable with primer G2 at the sgRNA target site (Fig. 2o and p, Fig. 2i and j). Again, no amplification was observed when using primers upstream and downstream of the sgRNA binding site (Fig. S2i and j). This indicates confined modification at the genomic region immediately proximal to the sgRNA recognition site and no extension to more distal positions within the assayed window. We confirmed NAc-specific plasmid expression in fresh coronal sections using GFP visualization and transfection efficiency via qPCR analysis 4 days post-surgery (Fig. S2k-m). These data established bidirectional epigenetic editing of *Cartpt* H3K27me3 and expression in the mouse NAc *in vivo*.

### Epigenetic editing of H3K27me3 in the nucleus accumbens regulates cocaine reward behavior

CART peptide is implicated in motivated behavior and drug addiction^19,28,42–45^. Our prior study found that *Cartpt* expression during cocaine abstinence is accompanied by loss of H3K27me3^11^. Therefore, we tested the hypothesis that *Cartpt* H3K27me3 is necessary and sufficient for *Cartpt* expression and cocaine reward behavior. We assessed cocaine conditioned place preference (CPP) and the extinction of acquired preference^11,46^, paradigms that align with the time-course of CRISPR-mediated epigenetic editing in NAc (Fig 3a). We used Social LEAP Estimates Animal Poses (SLEAP)^47^, a machine learning approach to automatically track freely moving mouse behavior and verified this with blinded manual scoring (Fig. 3b). We calculated preference scores for the cocaine and unrewarded chambers and set a threshold of acquired cocaine CPP when the acquisition delta preference score (calculated as preference score posttest minus pretest) was greater than 200 seconds.

**Figure 3.**
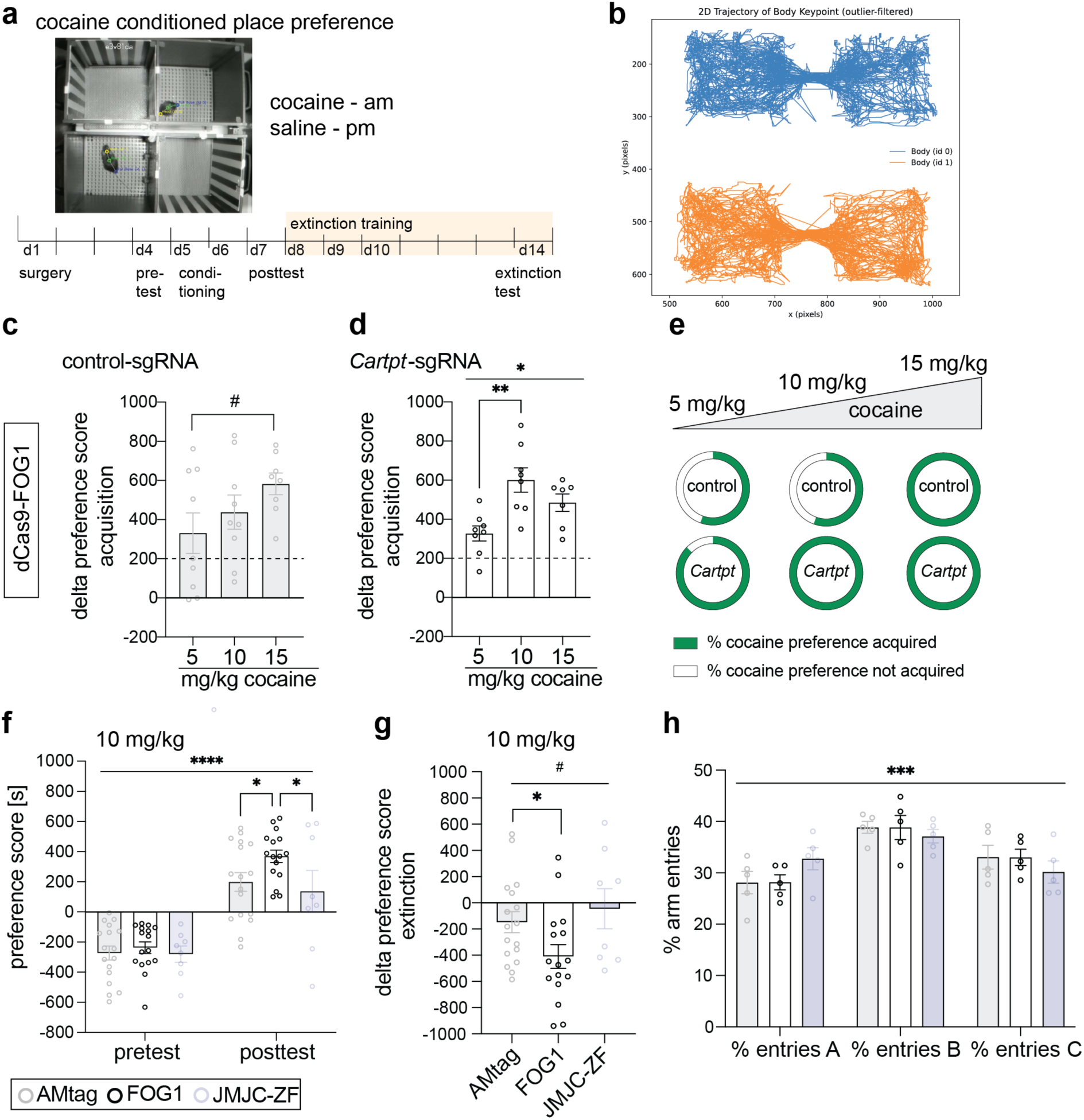
Epigenetic manipulation of *Cartpt* in the NAc regulates cocaine CPP acquisition and extinction. **a** Time course of CRISPR treatment, cocaine CPP and extinction. **b** Representative SLEAP analysis. **c** Dose response curve of cocaine CPP at 3 different doses, 5, 10, and 15 mg/kg cocaine. Control epigenetic editing with dCas9-FOG1 and control-sgRNA gradually increased cocaine preference leading up to 15 mg/kg. 15 mg/kg led to trending increase compared to 5 mg/kg. One-way ANOVA, effect of dose p=0.1489, Tukey’s multiple comparisons test p=0.3800 5 mg/kg vs 10 mg/kg, #p=0.0538 5 mg/kg vs 15 mg/kg, n=8-9. **d** Dose response curve of cocaine CPP at 3 different doses, 5, 10, and 15 mg/kg cocaine. Epigenetic editing with dCas9-FOG1 at *Cartpt* significantly increased cocaine preference at 10 mg/kg compared to 5 mg/kg, 15 mg/kg led to trending increase compared to 5 mg/kg. One-way ANOVA, effect of dose **p=0.0031, Tukey’s multiple comparisons test **p=0.0022 5 mg/kg vs 10 mg/kg, p=0.0960 5 mg/kg vs 15 mg/kg, p=0.2633 10 mg/kg vs 15 mg/kg, n=8. **e** Percent of animals that acquired cocaine preference at different doses of cocaine, 5 mg/kg: acquired_control-sgRNA_=55.6%, not acquired_control-sgRNA_=44.6%, acquired*_Cartpt_*_-sgRNA_=87.5%, not acquired*_Cartpt_*_-sgRNA_=12.5%. 10 mg/kg: acquired_control-sgRNA_=55.6%, not acquired_control-sgRNA_=44.6%, acquired*_Cartpt_*_-sgRNA_=100%, not acquired*_Cartpt_*_-sgRNA_=0%. 15 mg/kg: acquired_control-sgRNA_=100%, not acquired_control-sgRNA_=0%, acquired*_Cartpt_*_-sgRNA_=100%, not acquired*_Cartpt_*_-sgRNA_=0%, n number as described in Fig 3c and d. **f** dCas9-FOG1/*Cartpt-*sgRNA increased cocaine CPP (10 mg/kg cocaine) compared to dCas9-FOG1/control-sgRNA, while dCas9-JMJD-ZF had no effect on cocaine CPP. Two-way RM ANOVA, significant effect of test session, ****p<0.0001, n(dCas9-AMtag)=17, n(dCas9-FOG1)=16, n(dCas9-JMJC-ZF)=8. Followed by uncorrected Fisher’s LSD, posttest AMtag vs FOG1 *p=0.0290, posttest AMtag vs JMJC-ZF p=0.5067, posttest FOG1 vs JMJC-ZF p=0.0166, AMtag pretest vs posttest p<0.0001, FOG1 pretest vs posttest p<0.0001, JMJC-ZF pretest vs posttest p=0.0009. **g** Analysis of delta preference score (extinction minus posttest) of all three groups. dCas9-FOG1/*Cartpt*-sgRNA increased extinction compared to dCas9-AMtag/*Cartpt*-sgRNA. Brown-Forsythe and Welch’s ANOVA test, #p=0.0678 and 0.0702, multiple comparisons with unpaired t with Welch’s correction, AMtag vs FOG1 *p=0.0390, AMtag vs JMJC-ZF p=0.5634, FOG1 vs JMJC-ZF p=0.0633. **h** Analysis of spatial memory using Y-maze after treatment with dCas9-AMtag, dCas9-FOG1, and dCas9-JMJC-ZF all combined with *Cartpt*-sgRNA, measurement in % time spent in each arm (A, B, C). Two-way ANOVA, significant effect of ‘arm’ ***p=0.0004, no effect of ‘arm x treatment’, ‘treatment’, or ‘animal’, n=5 per group.

First, we compared the acquisition delta preference score of 3 independent cohorts of mice treated with CRISPR-dCas9-FOG1 control-sgRNA or *Cartpt*-sgRNA and conditioned with different doses of cocaine (5, 10, and 15 mg/kg). All doses produced a significant increase in preference for the cocaine-paired chamber relative to pretest (Fig. 3c and d). In the control animals, we observed a gradual increase in delta preference score with increasing cocaine doses (Fig. 3c). Control mice injected intra-NAc with dCas9-FOG1 and control-sgRNA acquired maximal CPP at 15 mg/kg cocaine, with all animals reaching the 200-second acquisition threshold (Fig. 3c). Contrary to this, mice injected intra-NAc with dCas9-FOG1 and *Cartpt*-sgRNA, acquired maximal CPP already at 10 mg/kg with all mice reaching the acquisition threshold (Fig. 3d and e, Fig. S3a). This indicates that H3K27me3-mediated *Cartpt* repression in NAc increased sensitivity to the rewarding effects of cocaine. In addition, intra-NAc infusions with dCas9-FOG1 and *Cartpt*-sgRNA increased the percentage of mice forming cocaine preference at 10 mg/kg by nearly 50% compared to non-targeting controls (Fig. 3e). A similar trend was observed at 5 mg/kg, whereas at 15 mg/kg, all mice acquired preference regardless of *Cartpt* H3K27me3 levels (Fig. 3e). Based on these findings, we selected a 10 mg/kg dose of cocaine for subsequent experiments and compared the effects of CRISPR-dCas9-FOG1 and dCas9-JMJC-ZF to control dCas9-AMtag. Mice treated with *Cartpt-*sgRNA and dCas9-FOG1 showed greater acquisition of cocaine CPP compared to dCas9-AMtag control (Fig. 3f). In contrast, animals treated with *Cartpt*-sgRNA and dCas9-JMJC-ZF showed no difference in acquisition of cocaine preference during the posttest compared to dCas9-AMtag control (Fig. 3f). Treatment with dCas9-FOG1 and dCas9-JMJC-ZF produced distinct behavioral outcomes, indicating that bidirectional editing of *Cartpt* H3K27me3 levels can yield differential effects, even when JMJC-ZF-mediated demethylation alone shows no behavioral change compared to controls (Fig. 3f). We note that unconditioned exploratory behavior, referred to as pretest, was not different between mice injected with *Cartpt-*sgRNA and dCas9-FOG1, dCas9-JMJC-ZF or dCas9-AMtag control. Further, there was a main effect of time, with pre- and posttest scores differing regardless of manipulation (Fig. 3f). Given the delayed activation of *Cartpt* during abstinence, we expanded the standard CPP protocol to include 4 extinction sessions over 1 week (Fig. 3a). We calculated an extinction delta preference score as the difference in time spent in the cocaine chamber during the posttest and the last extinction test on day 14. We found that dCas9-FOG1 with *Cartpt-*sgRNA increased extinction of cocaine preference compared to control (Fig. 3g, Fig. S3b). Despite CART peptide’s known effects on body weight^48^ we saw no effect of epigenetic editing on weight gain over 14 experimental days (Fig. S3c). Further, H3K27me3 and subsequently *Cartp*t levels had no impact on short-term working spatial memory during a modified Y-maze test^49^ test (Fig. 3h).

Taken together, we find that CRISPR-dCas9-FOG1 derepression of *Cartpt* during cocaine exposure stabilizes cocaine context memory across cocaine abstinence, supporting our hypothesis that *Cartpt* H3K27me3 is causally relevant to cocaine behavior. As there was no effect of CRISPR-dCas9-JMJC-ZF (supplementary Fig. 3a and b), we concluded that *Cartpt* H3K27me3 is sufficient but not necessary for cocaine reward behavior.

### Cell type-specific epigenetic editing of H3K27me3 regulates cocaine reward behavior

After discovering that enrichment of *Cartpt* H3K27me3 was sufficient to drive cocaine CPP and extinction (Fig. 3), we next investigated if this effect was mediated by either D1 or D2 MSNs, the two major neuronal subtypes in the NAc. First, we confirmed the cocaine-dependent expression of *Cartpt* in both D1 and D2 MSNs in a cell type-specific RNAseq dataset after 7 days of chronic cocaine exposure (Fig. 4a). To validate the cell type-specific vector, we previously performed cell type-specific CRISPRa targeting *Nr4a1,* a known upstream regulator of *Cartpt*^11^ in a separate series of experiments. To specifically target D1 or D2 NAc neurons, we injected Cre-dependent CRISPRa into the NAc of D2-Cre mice. We measured activation of the CRISPR target gene, *Nr4a1*, and *Nr4a1* target, *Cartpt*, only in the expected cell type (Fig. 4b and c). Next, we designed Cre-dependent dCas9-FOG1 and dCas9-JMJC-ZF for expression in D1 or D2 MSNs using D1-Cre or D2-Cre transgenic mice, respectively (Fig. S2a) and measured cocaine CPP as described above (Fig. 3). Based on D1 and D2-specific sequencing results where *Cartpt* was activated on both cell types by cocaine (Fig. 4a), we expected a behavioral effect in both cell type manipulations. We performed two separate sets of experiments with non-targeting sgRNA as control. Epigenetic repression of *Cartpt* in D1 MSNs increased cocaine CPP (Fig. 4d) and had no effect on extinction (Fig. 4e). *Cartpt* repression in D2 MSNs had no effect on either preference acquisition or extinction (Fig. 4f and g). This aligns with other studies that point towards a cell type-specific role of *Cartpt* in reward processing^31^. Based on these cell type-specific dCas9-FOG1 results, we next used Cre-mice as control to compare the effects of *Cartpt* activation in D1 and D2 MSNs in one experiment. Unexpectedly, we observed differences in pretest preference levels when activating *Cartpt* in D2 MSNs (Fig. 4h, see discussion). We observed no effect on CPP acquisition or extinction following cell type-specific H3K27me3 removal at *Cartpt* using dCas9-JMJC-ZF (Fig. 4h and i). Taken together, the absence of the above observed extinction effect with cell type-specific epigenetic editing suggests a cooperative role of H3K27me3 in D1 and D2 MSNs (Fig. 4j).

**Figure 4.**
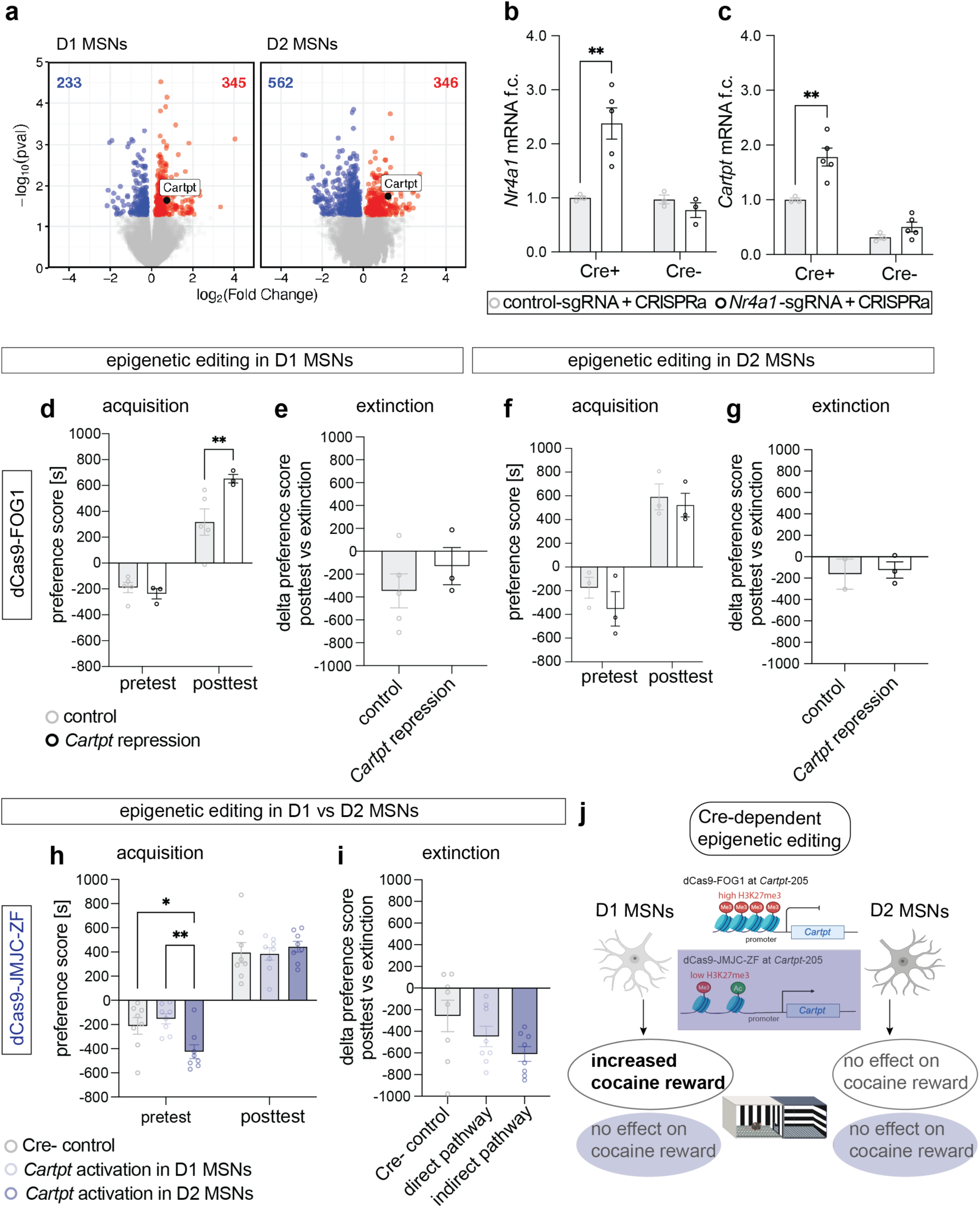
Cell type-specific manipulation of H3K27me3 at the *Cartpt* promoter regulates cocaine reward behavior via D1 MSNs. **a** Volcano plots of differentially expressed genes (DEGs) 1 day after 7 days of i.p. cocaine injections (20 mg/kg), individual DESeq2 objects were analyzed for each cell type using ‘treatment’ as a variable, left to right: DEGs in D1 MSNs, DEGs in D2 MSNs. Blue indicates significantly downregulated genes, red indicates significantly upregulated genes, padj<0.05, n=7-8. **b** Confirmation of Cre-specific epigenetic editing with *Nr4a1*-sgRNA and CRISPRa. *Nr4a1* mRNA was increased in *Nr4a1*-sgRNA transfected cells (black circles) relative to control-sgRNA (grey circles). Mixed-effects analysis, main effect genotype (p=0.001), treatment (p=0.1470), genotype x treatment (p=0.0012), Bonferroni’s multiple comparisons test: Cre+ control-sgRNA vs. Nr4a1-sgRNA **p=0.0062, Cre-control-sgRNA vs. Nr4a1-sgRNA p>0.9999, n=3-5. **c** Confirmation of Cre-specific epigenetic editing with *Nr4a1*-sgRNA and CRISPRa. *Cartpt* mRNA was increased in *Nr4a1*-sgRNA transfected cells (black circles) relative to control-sgRNA (grey circles). Mixed-effects analysis, main effect genotype (p<0.0001), treatment (p=0.0024), genotype x treatment (p=0.0368), Bonferroni’s multiple comparisons test: Cre+ control-sgRNA vs. Nr4a1-sgRNA **p=0.0018, Cre-control-sgRNA vs. Nr4a1-sgRNA p=0.6323, n=3-5. **d** D1 MSN-specific manipulation of H3K27me3 and *Cartpt* mRNA levels via dCas9-FOG1 and *Cartpt*-sgRNA increased cocaine preference compared to dCas9-FOG1 and control-sgRNA. All animals successfully acquired cocaine preference independent of treatment. Two-way ANOVA, followed by uncorrected Fisher’s LSD, pretest control- vs *Cartpt*-sgRNA p=0.6612, posttest control nt- vs *Cartpt*-sgRNA **p=0.0082, control pretest vs posttest p=0.0009, *Cartpt*-sgRNA pretest vs posttest p=0.0002, n=3-5. **e** D1 MSN-specific manipulation of H3K27me3 and *Cartpt* mRNA levels via dCas9-FOG1 and *Cartpt*-sgRNA had no effect on extinction of cocaine preference compared to dCas9-FOG1 and control-sgRNA, two-tailed Welch’s t test, p=0.3707, t=0.9828, df=5.021, n=3-5. **f** D2 MSN-specific manipulation of H3K27me3 and *Cartpt* mRNA levels via dCas9-FOG1 and *Cartpt*-sgRNA showed no difference in cocaine preference compared to dCas9-FOG1 and control-sgRNA. All animals successfully acquired cocaine preference independent of treatment. Two-way ANOVA, followed by uncorrected Fisher’s LSD, pretest control- vs *Cartpt*-sgRNA p=0.2962, posttest control- vs *Cartpt*-sgRNA p=0.6728, control pretest vs posttest p=0.0053, *Cartpt*-sgRNA pretest vs posttest p=0.0033, n=3. **g** Extinction was unaffected by manipulation of H3K27me3 and *Cartpt* mRNA levels via dCas9-FOG1 and *Cartpt*-sgRNA compared to dCas9-FOG1 and control-sgRNA, two-tailed Welch’s t test, p=0.8398, t=0.2367, df=1.593, n=2-3. **h** D1 and D2 MSN-specific manipulation of H3K27me3 and *Cartpt* mRNA levels via dCas9-JMJC-ZF and *Cartpt*-sgRNA had no effect on cocaine preference compared to Cre- control mice. All animals successfully acquired cocaine preference independent of treatment. Two-way RM ANOVA, followed by uncorrected Fisher’s LSD, effect of ‘test x genotype’ p=0.0197 and ‘test’ p<0.0001, n=8; pretest Cre- vs *Cartpt* activation in D1 MSNs p=0.4826, pretest Cre- vs *Cartpt* activation in D2 MSNs *p=0.0149, pretest *Cartpt* activation in D1 MSNs vs *Cartpt* activation in D2 MSNs **p=0.023, posttest Cre- vs *Cartpt* activation in D1 MSNs p=0.8957, posttest Cre- vs *Cartpt* activation in D2 MSNs p=0.5657, posttest *Cartpt* activation in D1 MSNs vs *Cartpt* activation in D2 MSNs p=4811, Cre-pretest vs posttest p<0.0001, *Cartpt* activation in D1 MSNs pretest vs posttest p<0.0001, *Cartpt* activation in D2 MSNs pretest vs posttest p<0.0001. **i** Extinction was unaffected by cell type-specific manipulation of H3K27me3 and *Cartpt* mRNA levels in D1 and D2 MSNs via dCas9-JMJC-ZF and *Cartpt*-sgRNA compared to Cre-mice, one-way ANOVA, p=0.0920, n=8. **j** Schematic of workflow of cell type-specific epigenetic editing and behavioral outcome.

## DISCUSSION

Chronic drug taking and abstinence permanently alter the way neuronal genes respond to drug exposure and drug-associated cues^3^. Such persistent alteration in gene responsivity likely promotes relapse and underlies the chronic nature of addiction^50^. Altered gene responsivity reflects persistent and global chromatin remodeling^9^. We aim to identify functionally relevant alterations to the epigenome at genes critical for the development of and/or recovery from chronic drug exposure and abstinence.

*Cartpt* is transcriptionally activated by psychostimulants, including cocaine^11,19,34,51^ and has repeatedly emerged in the context of cocaine addiction^44,48,52,53^. Human post-mortem studies show increased CARTPT mRNA in the NAc, replicated in rodents after acute cocaine exposure^6,19,20^. The relevance of *Cartpt* to cocaine addiction is further supported by studies of the encoded peptide, CART^18^, which demonstrates that intra-NAc infusion reduces cocaine-induced locomotion and reward^21–25^ suggesting a homeostatic role in drug-related behavior. While mechanisms remain unclear, CART responds to elevated extracellular dopamine and suppresses dopaminergic transmission^26–28^. Despite this evidence, cocaine regulation of *Cartpt* gene expression and chromatin architecture remains incompletely understood, and conflicting evidence across species and paradigms^54^ highlight the need to define cocaine-induced *Cartpt* expression in the NAc and assess its behavioral impact via gene-level manipulation.

Our current and prior studies find that *Cartpt* is repressed until several weeks of cocaine abstinence and identify a role for the repressive histone mark, H3K27me3, therein^11,34^. Integration of NAc gene expression data at 4, 7, 14 days of abstinence revealed that global transcriptional changes emerge primarily at 14 days, with minimal overlap between changes activated at earlier versus later stages of abstinence. Delayed *Cartpt* activation in mouse NAc reflects dynamic enrichment of repressive H3K27me3 across cocaine abstinence - enriched at 1 day and depleted at 28 days - and sustained enrichment of permissive histone modifications, H3K27ac and H3K4me3^11^. To determine the functional relevance of H3K27me3 to abstinence-delayed activation of *Cartpt,* we developed a novel, CRISPR-based epigenetic editing approach to manipulate *Cartpt* H3K27me3 in a single cell type in a single mouse NAc. We found that chromatin repression of *Cartpt* enhances acquisition and extinction of cocaine-associated memory. To the best of our knowledge, we are the first to apply targeted epigenetic editing of H3K27me3 in the brain.

We applied CRISPR-dCas9-FOG1 to specifically enrich H3K27me3 at *Cartpt.* FOG1 is a zinc finger protein that recruits the NuRD complex to enrich H3K27me3 and promote gene repression^35,37^. We modified this tool for neuronal-specific expression in mouse brain and showed successful enrichment of *Cartpt* H3K27me3 and reduction of *Cartpt* mRNA. For the removal of H3K27me3 and gene activation, we designed a novel tool based on the lysine-specific demethylase 6A (Kdm6a), containing a JMJC domain^40,41^. We tethered the JMJC effector domain and ZF domain to both the N- and C-termini of dCas9. We used only the JMJC catalytic domain to limit the demethylation capacity to the target locus, removing protein-protein interactions and minimizing off-target effects^55^. The JMJC domain catalyzes the removal of methylation groups from H3K27me3 and the ZF motif allows for the specific recognition of H3K27^40,41^.

Previous studies demonstrate that targeted epigenetic editing at addiction- and stress-related genes has successfully modulated behavioral responses to drugs and stress^56–58^ as well as cell type-specific memory expression^59^. The ability to reversibly alter gene expression without breaking DNA strands^60–62^ is a promising avenue for the development of targeted therapeutics for addiction. We present the first study on targeted epigenetic editing of H3K27me3 at the *Cartpt* promoter and prove its behavioral relevance in cocaine reward.

Using CRISPR-dCas9-FOG1 we showed that enrichment of *Cartpt* H3K27me3 is sufficient to repress *Cartpt* and drive the acquisition and extinction of cocaine preference. Further, as there was no effect of CRISPR-dCas9-JMJC-ZF, we concluded that *Cartpt* H3K27me3 is sufficient but not necessary for cocaine reward behavior. Global enrichment of permissive histone acetylation using small molecule histone deacetylase inhibitors selectively accelerates extinction without affecting initial acquisition of cocaine CPP^46^. Specifically, high-dose HDAC3 inhibitors enhance late extinction with no effect on acquisition^46^. These findings align with our observation that depletion of a repressive histone modification at *Cartpt* has limited effects on acquisition but may influence later stages of learning. Our findings support the hypothesis that reduced responsivity of the *Cartpt* gene increases drug context memory.

Due to the well-studied cell type-specific regulation of cocaine reward and extinction^31,63–68^, we applied our new CRISPR epigenetic editing tools in D1 and D2 MSNs in the NAc. We found that, unlike after enrichment of *Cartpt* H3K27me3 in all NAc neurons, enrichment of *Cartpt* H3K27me3 in D1 MSNs alone augmented acquisition but not extinction. This suggests that repression of *Cartpt* in both D1 and D2 MSNs is required to augment extinction. We note that mice expressing dCas9-JMJC-ZF in D2 MSNs showed baseline differences in chamber preference. However, this baseline preference was within the range of compartment preference across all cohorts and was present only in the Cre+ mice, indicating that it reflects the Cre+ transgene and not the epigenetic editing approach. Thus, while we acknowledge this baseline shift, we interpret it as genotype-related and unlikely to confound our conclusions regarding cocaine-induced preference.

Recent discoveries in neurobiological regulation of metabolism and feeding by similar peptides, such as GLP-1, emphasize the close relationship between innate responses to natural food rewards, and the overtaking of reward pathways by drugs^69^. GLP-1 agonists in the ventral tegmental area reduce cocaine self-administration in rats^70,71^ and show promise in rodent studies of opioid use disorder^72^. In parallel, efforts to identify a receptor for CART peptide remain ongoing^54,73–77^. Recent studies identify GPR-160 as a candidate receptor for CART peptide and implicate it in pain modulation^78^, meal microstructure^75^, and anorexigenic effects^73^. However, binding assay studies have complicated understanding GPR-160 as a functional receptor for CART^74,79^. CART peptide has also been shown to enhance memory formation and consolidation through modulatory effects on synaptic plasticity, neurotransmitter systems, and neurotrophic factors like BDNF. Cognitive-enhancing properties of CART make it a promising therapeutic target for memory-related disorders^76^, and is likely relevant to cocaine memory in our CPP assays.

Our results highlight the use of CRISPR-based epigenetic editing as a tool to investigate the direct causal relevance of gene-specific chromatin changes to behavior. By directly targeting H3K27me3 at the *Cartpt* promoter, we were able to induce specific transcriptional changes without altering the DNA sequence. This allows us to investigate the causal role of chromatin modifications in complex behaviors by targeting endogenous regulatory mechanisms instead of using overexpression or global manipulation of epigenetic modifiers. Our data supports epigenetic editing as a viable method to reprogram chromatin landscape relevant to highly prevalent psychiatric disorders. Future studies should assess downstream CART peptide levels to confirm transcriptional effects. Incorporating snRNA-seq and expanded CUT&RUN will help delineate mechanisms and exclude secondary effects on histone modifications and transcription. Extending behavioral analysis to a self-administration paradigm would further address compulsive drug-seeking behavior.

The present study extends our understanding of the brain’s response to cocaine. Our proposed mechanism shows that repression of *Cartpt* confirms its function as a regulatory brake on reward learning and shows a new role as a stabilizer of drug-seeking behavior. This could present the groundwork for new therapeutic strategies to address maladaptive memory formation and behaviors that underlie addiction and relapse.

## ACKNOWLEDGEMENTS

We thank Dr. Peter Hamilton for providing the hSyn-LS1L-GW-mCherry construct. We kindly acknowledge financial support from NIH/NIDA (R01DA052465 and U18DA052513 to E.A.H.). We would also like to thank the NIDA drug supply program for providing cocaine used in this study. The abstract schematic, Fig. 2d, d, f, m, and Fig. 4j were created using Biorender.com.

## AUTHOR CONTRIBUTIONS

J.J.W. and E.A.H. designed and analyzed behavioral experiments, SLEAP analysis and design was performed by R.W.D. Behavioral experiments and tissue collection were performed by J.J.W., A.A.G, K.S.C., and M.H. Animal husbandry, breeding, and genotyping was conducted by C.E.E. and K.S.C. RNA seq and CUT&RUN experiments were designed by J.J.W., E.A.H., and M.H., performed by J.J.W., analyzed by J.J.W. and K.S.K. Cell type-specific RNA seq experiments were designed by E.N. and B.W.H., performed and analyzed by B.W.H., M.E., C.D.T., A.R., and L.S. *In vivo* and *in vitr*o CRISPR studies wer designed by J.J.W. and E.A.H and performed by J.J.W., M.H., and K.L.R., construct design and cloning was performed by J.J.W., K.S.K, C.H., and K.L.R. qPCR and qChIP design and analysis was performed by J.J.W. and M.H. J.J.W. and E.A.H. wrote and revised the manuscript.

## DECLARATION OF INTEREST

The authors declare no competing interests.

## METHODS

### Animals

We used female and male 8-week old C57BL/6J mice (The Jackson Laboratory) in this study. Animals were housed under standard conditions on a 12-h light-dark cycle at a constant temperature of 23 °C and they had access to food and water ad libitum. For injections, animals were habituated to experimenter handling for 1 week before experimentation. For nuclear tagging for fluorescence-assisted nuclei sorting (FANS) followed by cell type-specific RNA-seq, transgenic D1-cre (Tg(Drd1-cre)FK150Gsat/Mmucd; GENSAT) and D2-cre (Tg(Drd2-cre)ER44Gsat/Mmucd; GENSAT) mice crossed to ROSA-Sun1/sfGFP (B6;129-Gt(ROSA)26Sor^tm5(CAG-Sun1/sfGFP)Nat/J; JAX 021039) were used. For cell type-specific epigenetic editing and behavioral analysis, R26-CAG-LSL-Sun1-sfGFP knock-in mice on the C57BL/6J background were crossed with A2a-cre and Drd1-cre mice C57BL/6J background to generate Sun1-sfGFP;A2a-cre and Sun1-sfGFP;Drd1-cre mouse lines. Guidelines of National Institutes of Health in Association for Assessment and Accreditation of Laboratory Animal care were followed for maintenance of all animals. All experiments and manipulations were approved by the Institutional Animal Care and Use Committee of The University of Pennsylvania.

### Investigator administered cocaine

For intraperitoneal (i.p.) injections mice underwent 3 handling sessions. Daily cocaine injections consisted of cocaine hydrochloride dissolved in sterile 0.9 % saline (20 mg/kg), control animals received a weight adjusted injection of sterile 0.9 % saline. Animals were allowed to move freely in an unknown cage for 15 minutes before being returned to their homecage. After last injection sessions, animals were kept in their homecage for 1, 4, 7, or 14 days of abstinence. Saline control animals underwent the corresponding abstinence regimen.

### Conditioned place preference

On day 1 of the protocol the mice underwent pretest and were allowed to freely explore the two-sided CPP chamber for 20 min to determine baseline preference. Animals that showed an initial preference of more than 70 % for one chamber were excluded from further analysis. Baseline preference score was calculated as non-preferred chamber minus preferred chamber in seconds. For conditioning animals received cocaine injection (5, 10, or 15 mg/kg) in the morning while in their non-preferred, now cocaine-paired, chamber for 30 min. In the afternoon, mice received a saline injection in their preferred chamber again for 30 min. To assess cocaine preference, mice were placed in the two-sided CPP chamber again and explored for 20 min. Preference score was calculated as time spent in cocaine-paired chamber minus time spent in saline-paired chamber in seconds. Delta preference score is represented as preference score pretest minus preference score posttest or preference score extinction minus preference score posttest. Extinction sessions were performed on 3 consecutive days after the initial posttest and followed the same protocol. After a 3-day recovery period the final extinction session was performed.

### Y-maze test

We performed Y-maze as previously described^49^ with minor modifications. In brief, mice were allowed to explore arms A and C of the maze for 15 minutes, arm B was closed off. After 4 hours in their homecage, mice were returned to the maze with access to arms A, B, and C for 10 minutes. Spontaneous alternations and % entries into each arm were analyzed.

### Behavioral analysis with SLEAP

We used the open source deep-learning based framework Social LEAP Estimates Animal Poses (SLEAP)^47^ version 1.4.1 via its graphical user interface (GUI) for multi-animal pose tracking to analyze behavior during the conditioned place perference. Unless indicated otherwise, we utilized the default parameters recommended by the GUI. First, we sampled and labeled 20 frames from 8 randomly selected videos (160 training frames) to construct a training data set. We chose to use three body points for tracking: head center, body center, and tailbase. After training, selected a video at random from each of our three cameras to analyze. We generated a new video with the network’s tracking estimates for each randomly selected video and visually confirmed tracking. After confirming tracking was satisfactory, we analyzed the remaining videos and saved the network estimates. To assess place preference, we divided the boxes into two regions with a 20 pixel boundary and calculated the amount of time the animal’s body was to the left or right of this boundary. Then we followed preference score calculations indicated in the section above (Cocaine conditioned place preference).

### Intra-NAc transfections

Bilateral intra-NAc infusions were performed as described previously^11,15,56^. We used the following coordinates relative to bregma at an angle of 10° from the midline to target the NAc: +1.6 anterior/posterior, +/-1.5 medial/lateral, -4.4 dorsal/ventral. For transfections, we used jetPEI (Polyplus Sartorius) according to the manufacturer’s instructions. We delivered plasmid DNA at a concentration of 1 µg/µl in 1.5 µl jetPEI/plasmid solution per side and animal at a flow rate of 0.2 µl per minute, followed by 5 min rest before removing the syringes (Hamilton). After surgery, animals were allowed to recover for 4-7 days and daily monitored.

### Epigenetic editing tool design and preparation

*Cartpt* guide RNA at -205 base pairs relative to TSS was designed using Benchling (Biology Software) and IDT CRISPR-Cas9 guide RNA design checker (Integrated DNA Technologies). All DNA fragments were purchased from IDT (Integrated DNA Technologies). All dCas9 epigenetic effector constructs were cloned into the hSyn-GW-IRES-mCherry backbone or into hSyn-LS1L-GW-IRES-mCherry vector for cell type-specific targeting. dCas9-FOG1 fusion was extracted from dCas9-FOG1[N+C]^35^ (Addgene #100085) by PCR using Q5 High-Fidelity DNA Polymerase (NEB) (primers in Supplementary Table 1) and cloned into hSyn-LS1L-GW-IRES-mCherry and hSyn-LS1L-GW-IRES-mCherry expression vectors using Gateway cloning (Invitrogen) according to the manufacturer’s instructions. For dCas9-JMJC-ZF fusion design, we obtained the jumonji C sequence followed by a zinc finger motif of the *Kdm6a* gene from the mouse mm10 reference genome and confirmed 96.37% sequence identity with the human (hg38) sequence. Using the same N- and C-terminal approach as FOG1, including the (GGS)^5^ amino acid linker sequences, we obtained a 249 amino acid long fragment that was synthesized by GenScript and inserted into the final expression constructs using the same Gateway cloning strategy as described above. dCas9-AMtag was obtained from Active Motif. All final constructs were analyzed by whole plasmid sequencing (Plasmidsaurus) and purified using EndoFree Plasmid Maxi Kit (Qiagen). As control non-targeting sgRNA was injected and treated as baseline for both molecular and behavioral analysis. For cell type-specific epigenetic editing, both control-sgRNA and targeting sgRNA in Cre-mice were used and treated interchangeably.

### Tissue collection, RNA extraction, and qRT-PCR

Mice were euthanized by cervical dislocation and brains were removed immediately. Brains were placed in ice-cold phosphate buffered saline containing a protease inhibitor cocktail (Roche cOmplete Protease inhibitor) and sliced into 1 millimeter thick sections on a cold tissue matrix. NAc was micro-dissected using a 2 mm or 1.2 mm diameter rapid-punch (WellTech). Investigator administered cocaine studies used 2 mm diameter sections for analysis, intra-NAc infusions with epigenetic editing CRISPR tools were confirmed using fluoresecent microscopy (Leica) to ensure correct targeting and visual plasmid expression. S3EQ cytoplasmic RNA isolation was performed as described previously^80^ using RNeasy Micro Kit (Qiagen), nuclei were immediately stored at -80°C or used for downstream qChIP or CUT&RUN. RT-PCR to generate cDNA was performed using iScript cDNA Synthesis Kit (Bio-Rad). For analysis of qRT-PCR, we used the ΔΔCt method. All primer sequences can be found in Supplementary Table 1.

### RNA sequencing and data analysis

RNA sequencing was performed at Azenta Life Sciences. RNA was isolated using the above described extraction protocol. To identify differentially expressed genes across cocaine abstinence, paired-end read quality was first assessed using FastQC (https://www.bioinformatics.babraham.ac.uk/projects/fastqc/), with read depth being greater than 45 million per sample and mean position quality score greater than 35 across all samples. After checking quality, reads were trimmed for all samples with BBTools bbduk function (https://jgi.doe.gov/data-and-tools/software-tools/bbtools/) using the included adapters.fa database to remove 3‘ sequencing adapters and low-quality bases (Q < 30; ktrim=r k=23 mink=11 hdist=1 qtrim=r trimq=30) from the 3‘ end. Trimmed reads were then pseudoaligned to the mm39 genome with GENCODE vM36 basic annotations and quantified using kallisto (v0.46.2)^81^ including a correction parameter for GC bias (--bias). The resulting quantifications were imported using the tximport package (v1.26.1)^82^ and normalized count data transformed using countsFromAbundance = “lengthScaledTPM”. Differentially expressed genes were identified between saline and cocaine treated animals at each abstinence timepoint (e.g., 4-day saline vs. 4-day cocaine) using DESeq2^83^ with log2(FoldChange) shrinkage calculated using apeglm^84^. Genes were considered significantly different between cocaine and saline with an adjusted pvalue < 0.05. Figures were generated using base ggplot2 v3.5.2^85^ (volcano plots, Figure 1), eulerr v7.0.2^86^ (Euler plots, Figure 1) and pheatmap v1.0.13^87^ (heatmap, Figure 1). Gene ontology enrichment analysis was performed using the enrichGO function from clusterProfiler v4.6.2^88^ with significant DEGs compared to a gene universe encompassing all genes expressed greater than 1 TPM in all samples. Ontology terms were taken from the “Biological Process” set and considered significant with a Benjami-Hochberg-corrected pvalue < 0.05.

### Cell type-specific RNA isolation and sequencing

NAc tissue was dounce-homogenized on ice in RNA-compatible lysis buffer (320 mM sucrose, 5 mM CaCl₂, 0.1 mM EDTA, 10 mM Tris-HCl pH 8.0, 1 mM DTT, 0.1% Triton X-100, plus RNase inhibitors). Homogenates were filtered (40 µm), layered over a 1.8 M sucrose cushion, and ultracentrifuged (∼71,000 × g at 4 °C for ∼1–1.25 h) to pellet nuclei. Pellets were resuspended gently in ice-cold PBS with RNase inhibitors. For FANS, nuclei were stained with DAPI and sorted for GFP⁺ Sun1 nuclei using a standard gating strategy (singlets → DAPI⁺ nuclei → GFP⁺). Drd1-cre;Sun1-GFP and A2a-cre;Sun1-GFP cohorts were processed in parallel. Sorted nuclei were collected directly into RNA lysis buffer. Total RNA was isolated with on-column DNase treatment. Low-input RNA-seq libraries were prepared using the RNA-specific kitSMARTer RNA-seq V3 kit (Takara) per manufacturer’s instructions. Libraries were sequenced on an Illumina platform.

### Chromatin immunoprecipitation

ChIP was performed on bilateral NAc 1 mm punches from 4-6 animals pooled or from bilateral, single-animal NAc 1.2 mm punches dissected as described above. Magnetic beads (Dynabeads M-280, Life Technologies) were bound to antibodies against either H3K27me3 (EMD Millipore 07-449) or H3K27ac (EMD Millipore 07-360). Nuclei were obtained using the above described S3EQ RNA isolation protocol. After fixation and chromatin extraction, chromatin was sheared using a Diagenode bioruptor XL at high intensity for 30 min (30 s on/30 s off) and incubated overnight with antibody-bound Dynabeads. The beads were then washed twice each with 1 ml of Low Salt Wash Buffer (20mM Tris, pH 8.0, 150mM NaCl, 2mM EDTA, 1% TritonX-100, 0.1% SDS), High Salt Wash Buffer (20mM Tris, pH 8.0, 500mM NaCl, 2 mM EDTA, 1% TritonX-100, 0.1% SDS), and TE Buffer (10mM Tris, pH 8.0, 1 mM EDTA). Finally, DNA was reverse cross-linked and purified (Qiagen Spin Column). qChIP primer sequences can be found in Supplementary Table 1.

### CUT&RUN

Nuclei for H3K27me3 analysis were obtained following the above describe S3EQ protocol. Bilateral 2mm NAc punches were used from single animals. Pellets were resuspended in Wash Buffer (20 mM HEPES-NaOH pH 7.5, 150 mM NaCl, 0.5 mM spermidine, 0.1% Triton X-100, 0.1% Tween-20, 0.1% BSA, protease inhibitors. CUTANA Concanavalin A Conjugated Paramagnetic Beads (Epicypher) were activated in binding buffer before they were added to the nuclei suspension. Nuclei and beads were rotated for 10 min at 4°C and then anti-H3K27me3 (Activ Motif, 39055, 1:50 dilution) antibody was added to the mix at 4°C over night with rotation. Bead-bound nuclei were washed 2x in digitonin buffer and CUTANA pAG-MNase (Epicyper) was added for 1h at 4°C with rotation to cut the DNA. Bead-bound nuclei were again washed 2x in digitonin buffer before targeted chromatin digestion and release. 100 mM CaCl_2_ was added for 30 min at 4°C with rotation before the reaction was stopped by adding stop buffer. Antibody-enriched fragments were released at 37°C for 10 min then 1 µl of 10% (wt/vol) SDS and 1.5 µl of proteinase K (20 mg/ml) was added followed by incubation at 50°C for 20 min. DNA clean up was performed using QiaQuick columns and buffers (Qiagen) according to the manufacturer’s instructions and DNA was eluted in 26 µl TE buffer. We used the NEB Ultra II library preparation kit and followed a previously optimized protocol^36^. Libraries were sequenced at Azenta Life Sciences.

### CUT&RUN data analysis

Demultiplexed files were provided by Azenta Life Sciences. Reads were trimmed with BBTools bbduk function (https://jgi.doe.gov/data-and-tools/software-tools/bbtools/) as described above and aligned to mm10 using bowtie 2 (v2.5.1)^89^. Alignment files were converted to BAM format, sorted, and indexed using SAMtools (v1.10)^90^. Duplicate reads were removed (Picard v3.0 MarkDuplicates (https://broadinstitute.github.io/picard/) or SAMtools v1.10 rmdup^90^). For visualization, de-duplicated BAMs were converted to bigWig with deepTools (v3.5.0)^91^ bamCoverage (--binSize 10 --normalizeUsing RPKM) and viewed in IGV (v2.12.2)^92^. Peaks were called with SICER2^93^ or SEACR^94^ and visualized in IGV.

### Neuro2a transfection

Neuro2a (N2a) cells (CCL-131, ATCC) were cultured cultured in EMEM with 10% heat inactivated fetal bovine serum (ATCC). Cells were sub-cultured by gentle trypsinization (trypsin/EDTA, ATCC) at 80% confluence. Potential mycoplasma contamination was assessed regularly. For transfection of plasmid DNA, cells were seeded at a density of 1 x 10^6^ cells in 12-well plates the day before treatment. Transfections occurred in Opti-MEM Reduced Serum Medium (Gibco) or serum-free growth medium. After 24 hours, RNA was isolated using the RNeasy Micro Kit (Qiagen) following the manufacturer’s instructions and qRT-PCR analysis was performed as described above.

### Statistics

Statistical tests were chosen based on the number of performed comparisons. Details on the specific test performed are described in the respective figure legends. Statistical significance was considered at p < 0.05, while p values greater than 0.05 and below 0.1 were considered trends. Data are presented as mean ± standard error of the mean (s.e.m). Outliers were identified as above or below mean 2x ± standard deviation. To ensure normal distribution, F tests of variance were conducted on all data sets. All experiments were replicated one to three times and data replication was confirmed. Analysis was performed using GraphPad Prism (Version 10.6.1), for macOS, GraphPad Software, Boston, Massachusetts USA, https://www.graphpad.com). Additionally, detailed information on each statistical test can be found in the source data file.

## Data availability

All source data used is available through GEO (will provided after publication). Constructs will be provided on Addgene after publication. All data are reported in main text and in supplementary data tables.

## Code availability

Transcriptome data is available on the Gene Expression Omnibus. We used published software for data analysis and we can provide more detailed information by request to the corresponding author. Conditioned place preference behavioral analysis using SLEAP, code was written by RWD and is available upon request via the corresponding author.

## Notes

### Competing Interest Statement

The authors have declared no competing interest.

