## extended data for "Epigenetic editing of Cartpt promotes acquisition and extinction of cocaine memory"

Elizabeth A. Heller, PhD

10-115 Smilow Center for Translational Research

3400 Civic Center Boulevard, Building 421

Philadelphia, PA 19104-5158

**Supplementary Figure S2.**

**a** CRISPR plasmid maps

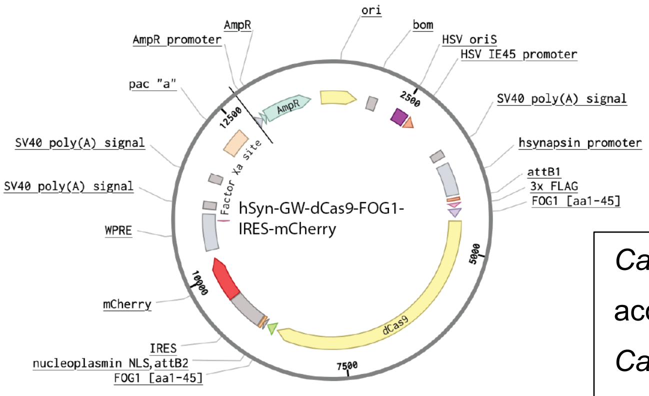


Cartpt-sgRNA sequence TSS -205:

accggacgtggtaccagtcc

*Cartpt*-sgRNA sequence TSS -526:

tattgtggcgagggtaacta

control-sgRNA:

cgaggtattcggctccgcg

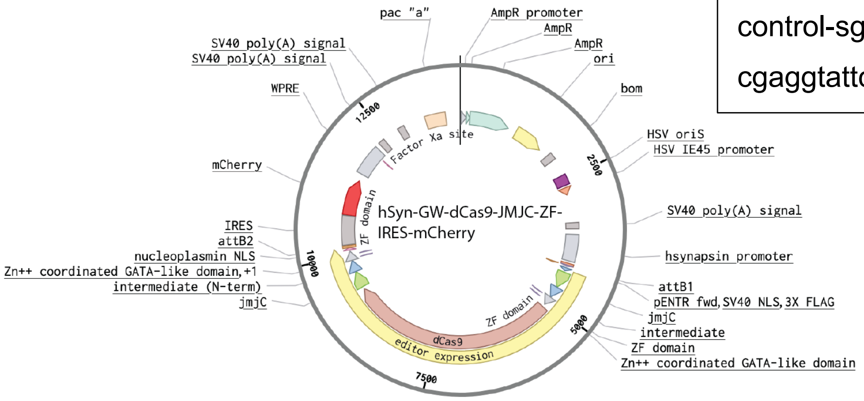


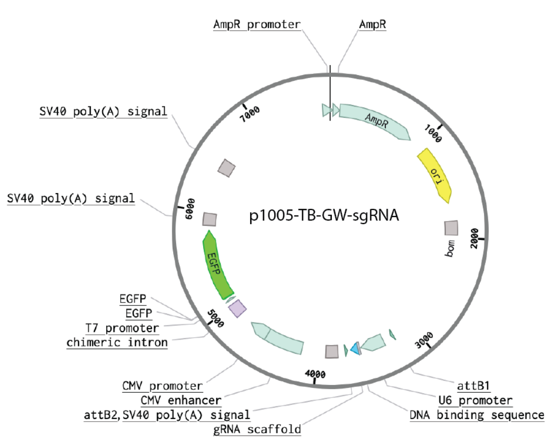


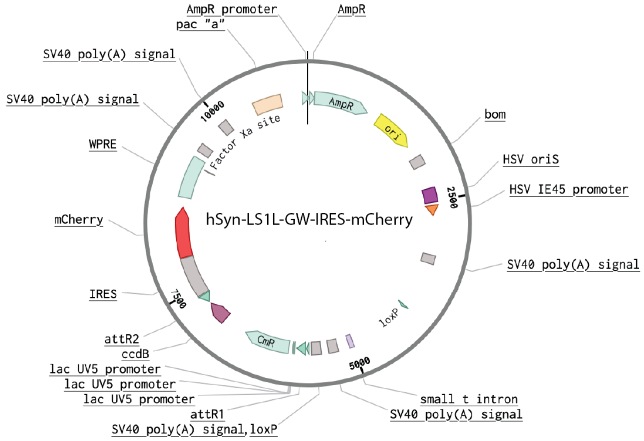

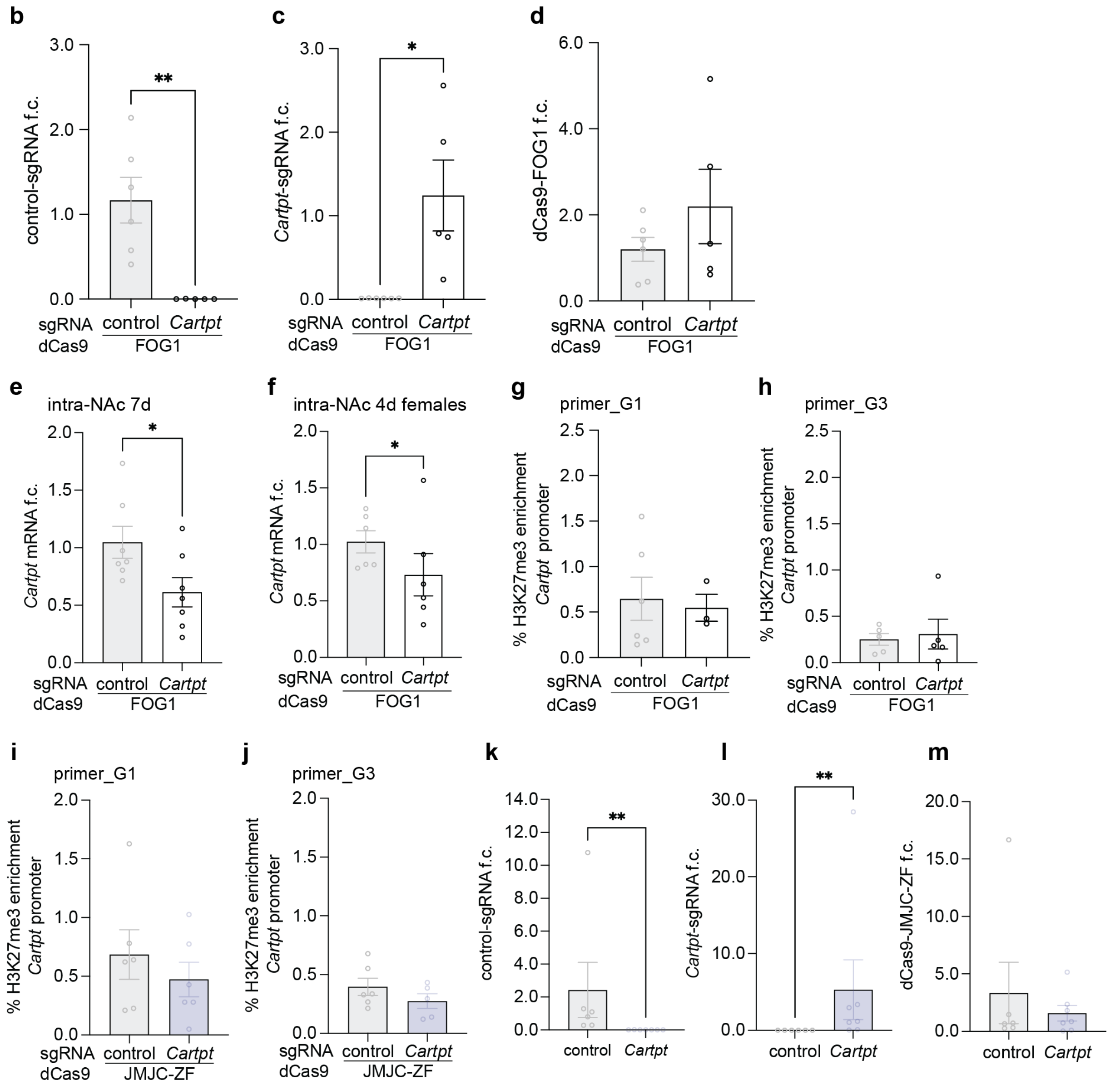


**Figure S2. Control qPCRs of epigenetic editing and additional qChIP analysis.**

**a** Plasmid maps of all constructs used in this study, generated with Benchling.com. Single-guide RNA sequences are as followed: *Cartpt*-sgRNA sequence TSS -205 accggacgtggtaccagtcc; *Cartpt*-sgRNA sequence TSS -526 tattgtggcgagggtaacta; control sg-RNA sequence cgaggtattcggctccgcg. **b** Control qPCR analysis for correct plasmid expression confirmed detectability of control-sgRNA in control samples with dCas9-FOG1, two-tailed t test, Mann-Whitney U=0, **p=0.0043, n=5-6. **c** *Cartpt*-sgRNA was only detectable in targeting samples and dCas9-FOG1 via qPCR, unpaired two-tailed t test with Welch's correction *p=0.0442, t=2.897, df=4.000, n=5-6. **d** Control qPCR confirmed equal dCas9-FOG1 expression in control and *Cartpt*-

sgRNA samples, unpaired two-tailed t test with Welch's correction p=0.3245, t=1.097, df=4.823, n=5-6. **e** *Cartpt* mRNA remained downregulated in NAc at day 7 after injections, compared to control-sgRNA, unpaired two-tailed t-test with Welch's correction, *p=0.0398, t=2.308, df=11.91, n=7. **f** *Cartpt* mRNA was downregulated in adult female mice treated with *Cartpt*-sgRNA and dCas9-FOG1 on experimental day 4 compared to control-sgRNA, paired one-tailed t test *p=0.0453, t=2.092, df=5, n=6. **g** Enrichment of H3K27me3 at the *Cartpt* promoter by dCas9-FOG1/*Cartpt*-sgRNA was not detectable when assessed by primer_G1, paired t test, p=0.4576, t=0.9131, df=2, n=3-6, relative to dCas9-FOG1 and control-sgRNA. **h** Enrichment of H3K27me3 at the *Cartpt* promoter by dCas9-FOG1/*Cartpt*-sgRNA was not detectable when assessed by primer_G3, Wilcoxon matched-pairs signed rank test, two tailed, p>0.9999, n=5, relative to dCas9-FOG1/control-sgRNA. **i** De-enrichment of H3K27me3 at the *Cartpt* promoter by dCas9-JMJC-ZF/*Cartpt*-sgRNA was not detectable when assessed by primer_G1, unpaired t test with Welch's correction, p=0.2164, t=0.8213, df=8.937, n=6, relative to dCas9-JMJC-ZF/control-sgRNA. **j** De-enrichment of H3K27me3 at the *Cartpt* promoter by dCas9-JMJC-ZF/*Cartpt*-sgRNA was not detectable when assessed by primer_G3, unpaired t test with Welch's correction, p=0.1207, t=1.254, df=8.988, n=5-6, relative to dCas9-JMJC-ZF/control-sgRNA. **k** Control qPCR analysis for correct plasmid expression confirmed detectability of control-sgRNA in control samples with dCas9-JMJC-ZF, two-tailed t test, Mann-Whitney U=0, **p=0.0012, n=6-7. **l** *Cartpt*-sgRNA was only detectable in targeting samples and dCas9-JMJC-ZF via qPCR, two tailed t test, Mann-Whitney U=0, **p=0.0012, n=6-7. **m** Control qPCR confirmed equal dCas9-JMJC-ZF expression in control and *Cartpt*-sgRNA samples, two tailed t test, Mann-Whitney U=19, p=0.8357, n=6-7.

**Supplementary Figure S3.**

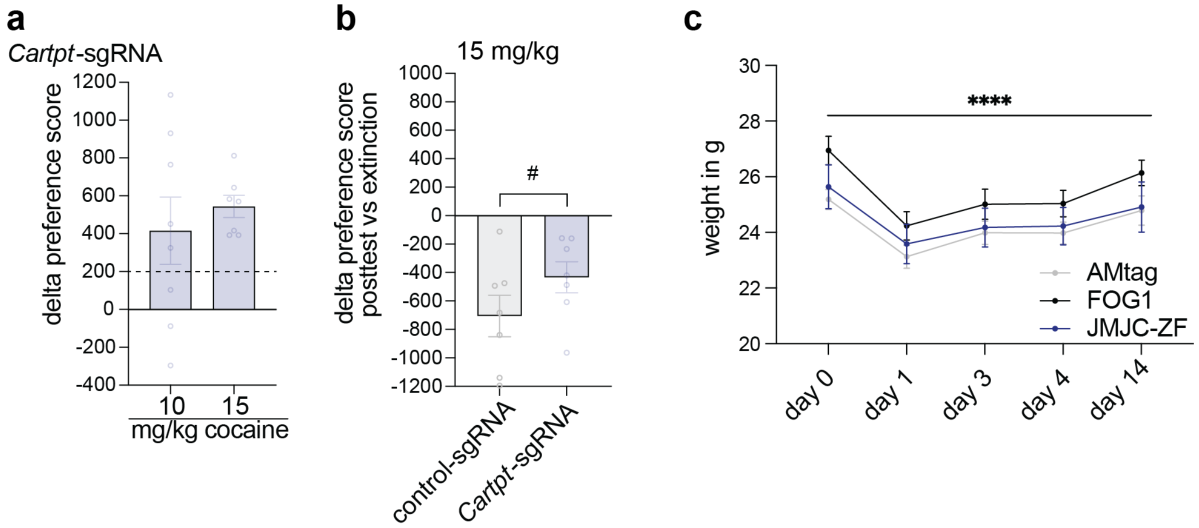


**Figure S3. Additional cocaine CPP data after epigenetic editing with dCas9-JMJC-ZF.**

**a** Dose response curve of cocaine CPP at 2 different doses, 10, and 15 mg/kg cocaine. *Cartpt*-sgRNA: epigenetic editing with dCas9-JMJC-ZF at *Cartpt* narrows cocaine preference at 15 mg/kg compared to 10 mg/kg, unpaired t test with Welch's correction, p=0.2552, one-tailed, t=0.6871, df=8.510. **b** Analysis of delta preference score posttest vs extinction of CPP at 15 mg/kg cocaine. Extinction of mice treated with dCas9-JMJC-ZF and *Cartpt*-sgRNA was decreased compared to control-sgRNA, unpaired t test with Welch's correction, #p=0.0821, one-tailed, t=1.489, df=11.12. **c** Tracking of body weight over 14 experimental days, two-way ANOVA, significant effect of time ****p<0.0001 and animal p<0.0001, no effect of treatment p=0.2727, n=8.

**Supplementary Table S1.** Primer design.

| Primer | Sequence |
|---|---|
| *Cartpt*_ori | fwd: TACTCTGCCGTGGATGATGCGT<br>rev: TCGGAATGCGTTTACTCTTGAGC |
| *Cartpt*_MDC (Carpenter et al., 2020) | fwd: ACGAGAAGGAGCTGATCGAA<br>rev: TCTCTGAGGGGAACGCAAAC |
| G1_primer | fwd: GGGTGTAGCAGCAAGAAGGA<br>rev: GTACCACGTCCGGTTCTCTC |
| G2_primer | fwd: CTGAACGGGAGCGAGAGAAC<br>rev: TGTGTACACGAGTGCAGGTG |
| G3_primer (Carpenter et al., 2020) | fwd: ACACAAGAGCCGTCAATTCCA<br>rev: TCGAGTTCCCAACACCGC |
| pENTR | fwd: CACCATGCCCAAGAAGAAGAGGAAG<br>rev: CTTTCATGCGGCCGCAGGTC |
| dCas9-FOG1 | fwd: CAGAGAGGAGGTGCAGTTGG<br>rev: CCGTGCTGTTCTTTTGAGCC |
| dCas9-JMJC-ZF | fwd: ACGGTGGTAGTGGAGGTTCA<br>rev: AAGGTTGGGCCACCAAGAAC |
| nt control-sgRNA | fwd: GCGAGGTATTCGGCTCCGCG<br>rev: TAATGCCAACTTTGTACAAGAAAG |
| *Cartpt*-sgRNA | fwd: ACCGGACGTGGTACCAGTCC<br>rev: TAATGCCAACTTTGTACAAGAAAG |
